# Peripheral, intrareceptor inhibition in mosquito olfaction

**DOI:** 10.1101/243162

**Authors:** Pingxi Xu, Young-Moo Choo, Zhou Chen, Fangfang Zeng, Kaiming Tan, Tsung-Yu Chen, Anthony J. Cornel, Nannan Liu, Walter S. Leal

## Abstract

How chemical signals are integrated at the peripheral sensory system of insects is still an enigma. Here we show that when coexpressed with Orco in *Xenopus* oocytes, an odorant receptor from the southern house mosquito, CquiOR32, generated inward (regular) currents when challenged with cyclohexanone and methyl salicylate, whereas eucalyptol and fenchone elicited inhibitory (upward) currents. Responses of CquiOR32-CquiOrco-expressing oocytes to odorants were reduced in a dose-dependent fashion by coapplication of inhibitors. This intrareceptor inhibition was also manifested in vivo in fruit flies expressing the mosquito receptor CquiOR32, as well in neurons on the antennae of the southern house mosquito. Likewise, an orthologue from the yellow fever mosquito, AaegOR71, showed intrareceptor inhibition in the *Xenopus* oocyte recording system and corresponding inhibition in antennal neurons. Intrareceptor inhibition was also manifested in mosquito behavior. Blood-seeking females were repelled by methyl salicylate, but repellence was significantly reduced when methyl salicylate was coapplied with eucalyptol.

**One Sentence Summary:** Intrareceptor inhibition was observed in mosquito odorant receptors expressed in heterologous systems, in vivo, and manifested in behavioral responses.

## INTRODUCTION

Integration of chemical signals at the peripheral sensory system (antennae, maxillary palps, and proboscis) remains one of the least understood mechanisms of insect olfaction, particularly in mosquitoes. Despite the great progress made in the last 2 decades in understanding how receptors form the basis of chemosensory perception in insects, how olfactory (as well as taste) signals integrate at the periphery remains an enigma (*1*). “It is as if a new continent has been discovered but only the coastline has been mapped” (*1*). In the largest majority of reported cases *(2-5)*, antennal neurons of *Cx. quinquefasciatus* displayed excitatory responses (increased spike frequency upon stimulus), but evidence for inhibitory responses (reduction in spontaneous activity upon stimulus), already known for *Ae. aegypti* (*6*), is now emerging for *Cx. quinquefasciatus* (*5*). It has been observed in moths (*7*), beetles (*8*), the fruit fly (*9*) and mosquitoes (*10*) that activation (firing) of one neuron interferes with signaling of other olfactory receptor neurons (ORNs; also referred to as olfactory sensory neurons, OSNs). It has also been reported that a single compound can elicit excitatory and inhibitory responses from the same neuron (*11*). Although Carlson and collaborators elegantly demonstrated that in the fruit fly lateral inhibition is most likely mediated by ephaptic coupling (*9*), the complete ensemble of the molecular mechanism(s) of inhibition at the peripheral olfactory system of mosquitoes remains *terra incognita*. A simple explanation of the ephaptic coupling is that upon (continuous) stimulation of an ORN the (external) potential (of the sensillum lymph surrounding dendrites) declines. Consequently, per channel current generated by a cocompartmentalized neuron (when stimulated by its cognate ligand) is reduced (*12*). This scenario argues that the firing of a neuron causes reduced spike frequency by a colocated neuron due to the close apposition (ephaptic, Greek for “to touch”) of their neuronal processes. Although ephaptic coupling could explain lateral inhibition, other mechanisms of intraneuron inhibition may exist. While de-orphanizing odorant receptors (ORs) expressed predominantly in *Cx. quinquefasciatus* female antennae, we serendipitously recorded currents that generate inhibition in response to certain odorants. Further studies unraveled a hitherto unknown mechanism of peripheral, intrareceptor inhibition in mosquito olfaction.

## RESULTS AND DISCUSSION

### Recordings of currents in upward direction

In our attempts to de-orphanize ORs from the southern house mosquito, *Cx. quinquefasciatus*, we challenged *Xenopus* oocytes coexpressing CquiOR32 along with the obligatory coreceptor Orco with a panel of ~250 physiologically and behaviorally relevant compounds *(13)*. Because CquiOR32 is predominantly expressed in female antennae (fig. S1), we reasoned that this receptor might be involved in the reception of attractants or repellents and therefore be of value to understanding critically important chemoreceptive egg laying and blood feeding behaviors of mosquitoes. CquiOR32-CquiOrco-expressing oocytes generated dose-dependent inward (regular) currents when challenged with various odorants, including cyclohexanone, methyl salicylate, and 2-methyl-2-thiazoline (Fig. 1A). Interestingly, however, the egg-laying deterrent, eucalyptol *(14)*, and many other compounds, including, fenchone, DEET, picaridin, IR3535, and PMD, generated currents in reverse direction (Fig. 1A). These unusual currents of reverse direction were reproducible, and no indication was found of adaptation (Fig. 1A, inset). No currents were recorded when oocytes alone or oocytes expressing only CquiOR32 or only CquiOrco were challenged either with methyl salicylate or eucalyptol (Fig. 1B). However, CquiOR32-CquiOrco-expressing oocytes responded in a dose-dependent manner to both compounds eliciting inward currents (cyclohexanone and methyl salicylate) (Figs. 1B, fig. S2) and those eliciting currents in the reverse direction (inhibitors; eg, eucalyptol, Fig. 1B, fig. S3).

**Fig. 1.**
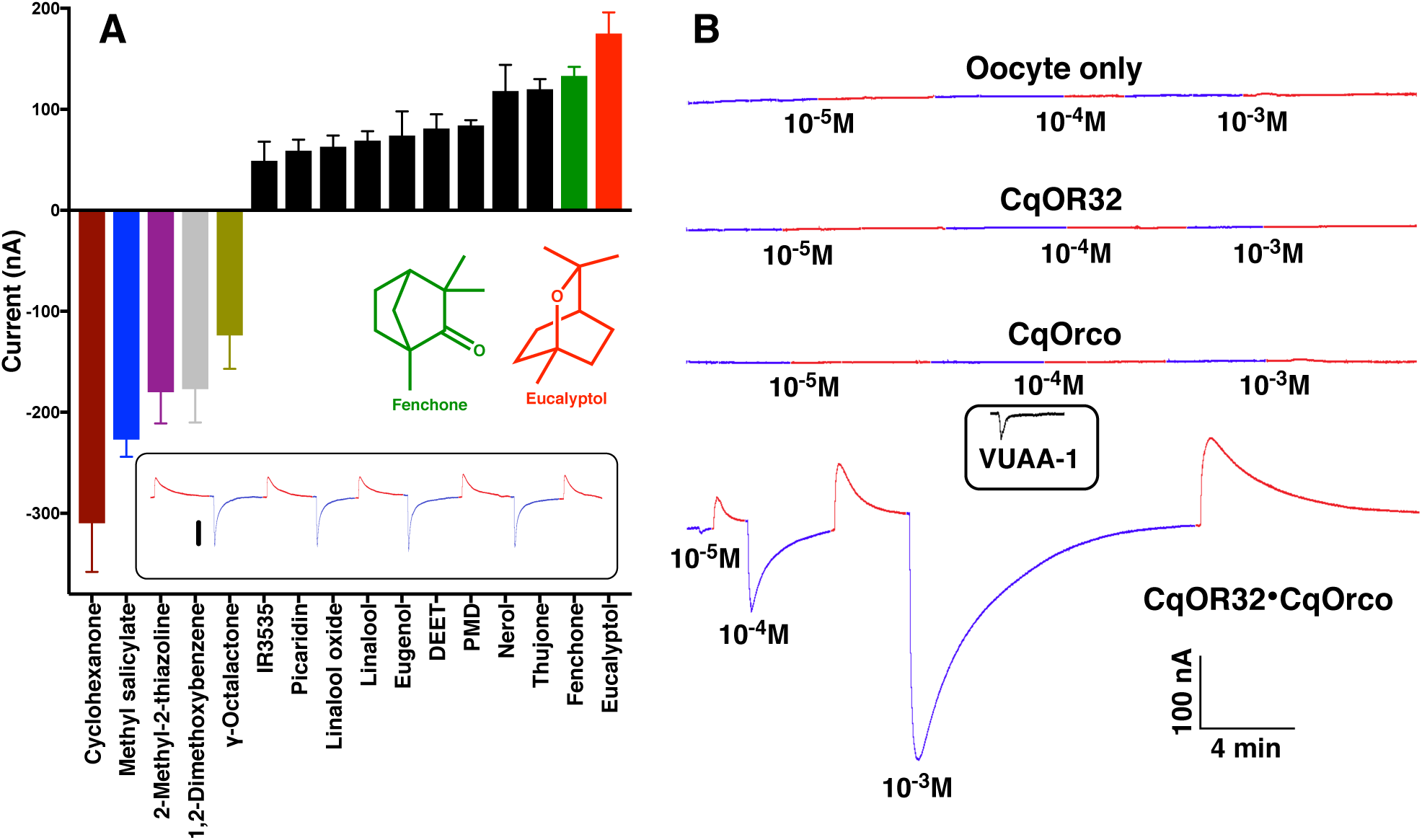
Recordings and quantification of inward and inhibitory currents. (A) CquiOR32-CquiOrco-expressing oocytes responses to ligands at 1 mM. Error bars represent SEM (n = 3–4). For clarity, bars representing reverse and inward currents are displayed upward and downward, respectively. The following compounds are not displayed for conciseness. Inward current-eliciting compounds: γ-hexalactone, 115±14 nA (all data in absolute values); guaiacol, 101±11 nA; acetophenone, 58±12 nA; 2-butanone, 45±7 nA; 2-phenylethanol, 44±9 nA. Inhibitory compounds: α-terpinene, 43±5 nA; terpinolene, 38±8 nA; citral, 37±17 nA; ocimene, 36±19 nA; geranylacetone, 30±5 nA; and α-pinene, 26±12 nA. (B) Control experiments with oocytes only and oocytes expressing only CquiOR32, CquiOrco, or a combination of CquiOR32 and CquiOrco. Response of a CquiOrco-expressing oocyte to Orco agonist, VUAA-1, is displayed in an inset. *Inset* in A: Continuous recordings from a single oocyte expressing CquiOR32-CquiOrco and repeatedly stimulated with 1 mM eucalyptol (reverse peaks; traces in maraschino) and 0.1 mM methyl salicylate (inward peaks; traces in blueberry).

### Current-voltage curves

To gain better insights into possible mechanism(s) of these currents in reverse direction, we obtained current-voltage (I-V) curves from oocytes expressing wild type (WT) CquiOR32-CquiOrco with voltage clamped at −80, −60, −40, −20, 0, +20, and +40 mV and using methyl salicylate and eucalyptol as stimuli (Fig. 2A&D, respectively). Similar recordings were performed using N-methyl-D-glucamine chloride (NMG) as a source of bulky, impermeable mono cation (Fig. 2B,E) or with sodium gluconate buffer (a source of bulky, impermeable anions) instead of NaCl (Fig. 2C,F). These data corroborate that OR32-Orco forms nonselective cation channels *(15–17)*, as indicated by the reversal potential shift to more negative voltage upon replacement of Na+ by less permeable NMG (Fig. 2A&B). An examination of the first and third quadrants for each graphic in Fig. 2 shows that I-V curves elicited by methyl salicylate (Fig. 2A-C) are above baseline and below control I-V curves for positive and negative voltages, respectively (a common pattern of inward currents), consistent with a potentiation of the control current by methyl salicylate. By contrast, I-V curves generated with eucalyptol, albeit not robust, are below and above control I-V curves for positive and negative holding potentials, respectively (Fig. 2D,E), suggesting an inhibition effect. In the presence of eucalyptol, replacing Na^+^ with NMG or Cl^-^ by gluconate had a minor effect on the I-V curves (Fig. 2E&F).

**Fig. 2.**
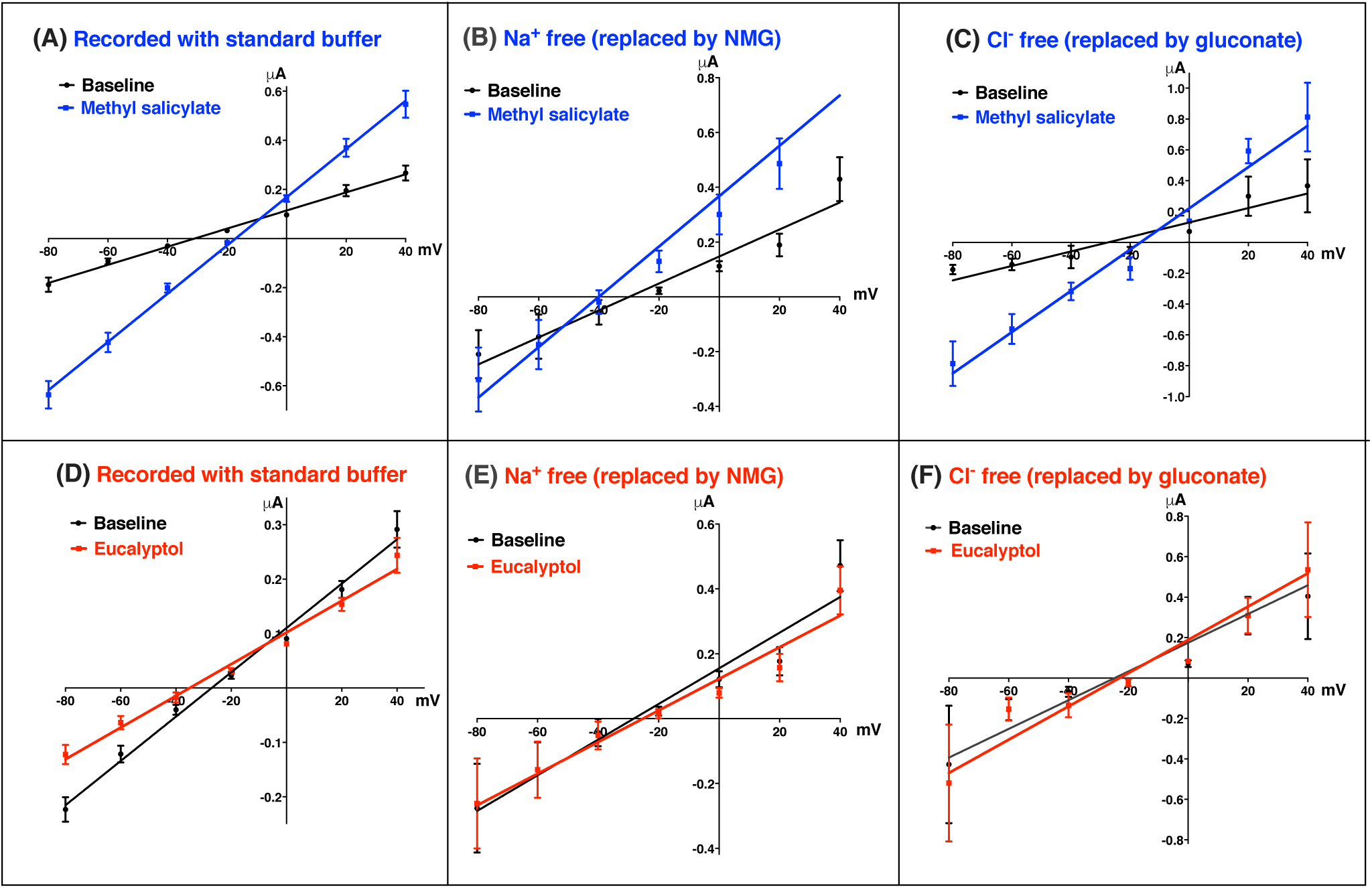
Current-voltage (I-V) curves for CquiOR32-CquiOrco expressed in *Xenopus* oocytes. Recordings with methyl salicylate (blueberry) and eucalyptol (maraschino) at 1 mM, with voltage clamped at −80, −60, −40, −20, 0, +20, and +40 mV and using regular and modified buffers. (A & D, n = 9) standard buffer; (B & E, n = 5) Na^+^ free buffer; (C&F, n = 3) Cl^-^ free buffer. Error bars represent SEM.

In the mammalian olfactory system, Cl^-^ efflux (inward current) depolarizes the membrane further *(18)*, thus providing the most pronounced amplification of the odor signal within the whole signal transduction cascade *(19)*. Cl^-^ influx (outward current) has been reported in taste cells of the common mudpuppy, *Necturus maculosus (20, 21)*. In insects, the difference in chemical composition of the sensillum (=receptor) lymph and hemolymph generates the standing electrical potential, ie, the transepithelial potential and it favors a possible Cl^-^ influx (*22*): sensillum lymph, K^+^200 mM; Na^+^ 25 mM; Cl^-^, 25 mM; hemolymph, K^+^, 36 mM; Na^+^ 12 mM; Cl^-^, 12mM; organic anions make the balance) (*23*).

It is worth mentioning that *Xenopus* oocytes have 2 types of endogenous Ca^2+^-activated Cl channels, ie, I_Cl-1_ and I_Cl-2_, which are localized in the animal hemisphere of the oocyte (*24*), but a few lines of evidence strongly suggest that our recorded inhibitory currents are not derived from oocytes. First and foremost, these currents are not generated by oocytes alone, or oocytes expressing only CquiOrco or CquiOR32; coexpression of CquiOR32 and CquiOrco is essential (Fig. 1). Additionally, *Xenopus* outward currents occur only when oocytes are depolarized to a potential more positive than −20 mV to cause release of Ca^2+^ from internal stores and activate I_Cl−1_; this outward current (due to Cl^-^ influx) has a time-to-peak of 200 ms, and is immediately followed by an inward current due to Ca^2+^ influx that activates and causes Cl^-^ efflux (*24-27*). By contrast, our newly recorded inhibitory currents are dose-dependent, and are never followed by inward currents. Lastly, currents recorded with Ca^2+^-free buffer (replaced by Ba^2+^) showed no significant difference (fig. S4).

We then surmised that these “inhibitors” might modulate responses to odorants when these 2 types of stimuli are simultaneously delivered to receptors. For example, the inward currents generated by methyl salicylate were outweighed by eucalyptol-induced reverse currents, thus generating a net (overall) current in the upward direction. As the doses of eucalyptol were gradually decreased, the reverse signal was reduced, cancelled out, and “overall” inward currents appeared (fig. S5), thus suggesting that “intrareceptor” inhibition occurred. Consistent with these findings, recordings of outward currents from antennal neurons (*28*) of the fruit fly, *Drosophila melanogaster*, have just been reported for the first time.

### Inhibition in the antennae of flies expressing CquiOR32

To test whether the inhibitory responses were manifested in vivo at the periphery of the olfactory system, we generated transgenic flies, with CquiOR32 expression driven by DmelOrco promoter, and recorded EAG responses by using a standard method (*29*). Control flies did not respond to eucalyptol, but gave basal responses only to higher doses of methyl salicylate. By contrast, CquiOR32 flies generated robust EAG responses when stimulated with methyl salicylate. As expected, 2-heptanone, the best EAG ligand for the fruit fly (*30*), elicited the greatest EAG responses. We did not observe significant EAG responses from CquiOR32 flies when challenged with eucalyptol per se. However, when delivered simultaneously through 2 separate cartridges, eucalyptol attenuated responses elicited by both 2-heptanone and methyl salicylate in a dose-dependent manner (Fig. 3). In control flies (UAS-CquiOR32) eucalyptol did not interfere with the normal responses to 2-heptanone (Fig. 3, inset). This dataset suggests that intrareceptor inhibition works (*31*) in vivo as indicated by the effect of eucalyptol on the methyl salicylate responses. A possible explanation for the reduction of 2-heptanone responses elicited by eucalyptol in a dose-dependent manner is that intraneuronal inhibition occurred, ie, CquiOR32 interacted with *Drosophila* innate OR(s) (eg, Or33a, 49a, 85b) (*31*).

**Fig. 3.**
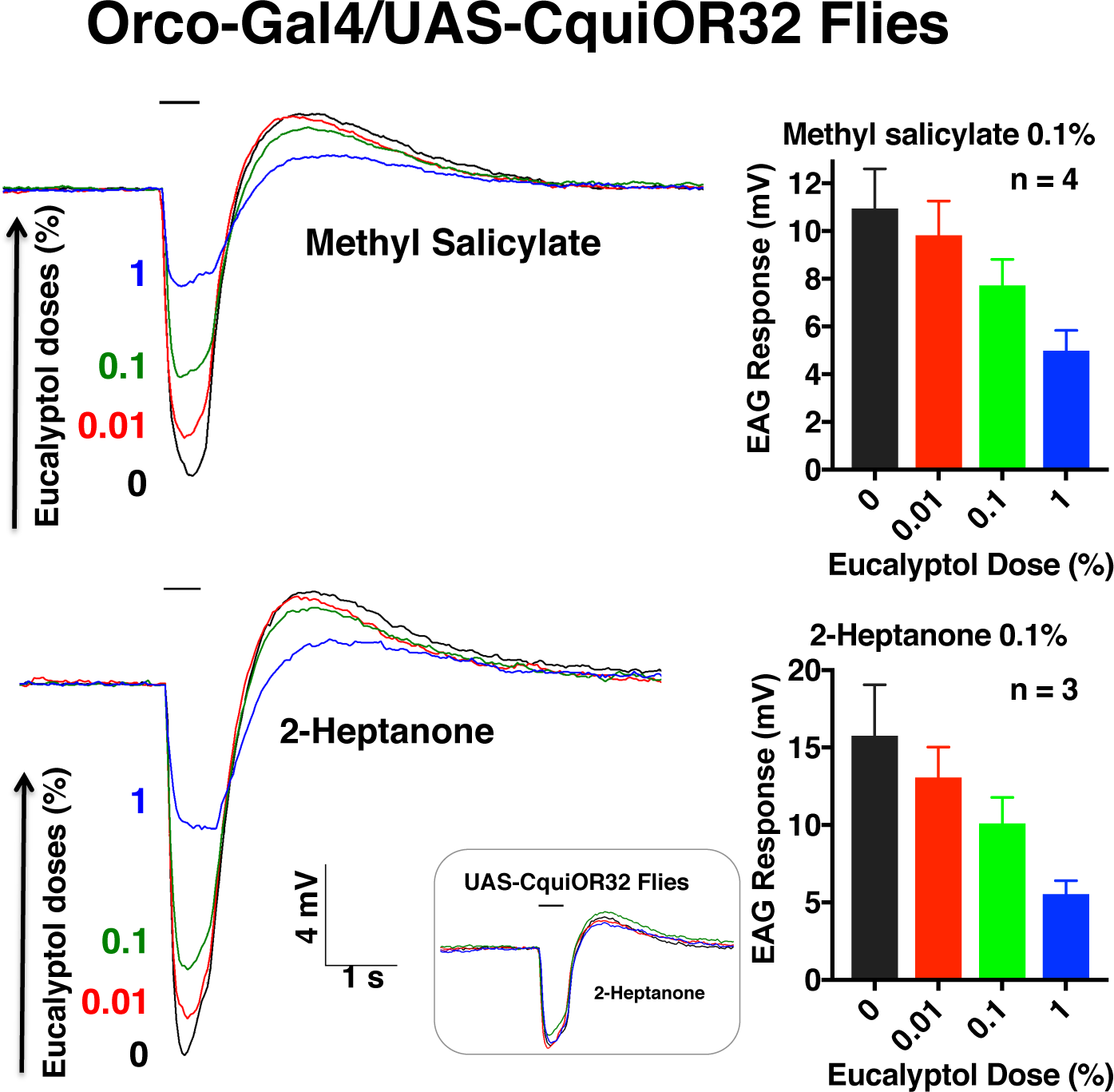
Inhibitory effect of eucalyptol on odorant reception in transgenic flies. Odorant (0.1%) was codelivered onto fly antennae with eucalyptol at various doses. Error bars represent SEM. *Inset:* EAG recordings from a control line.

### Inhibition in *Culex* mosquito antennae

We investigated by using single-sensillum recordings (SSR) whether inhibition would occur in antennae of the southern house mosquito. ORN-A in short sharp-tipped sensilla 2 (SST-2) responded to cyclohexanone with dose-dependent excitatory responses (Fig. 4A) and to eucalyptol (Fig. 4B) and fenchone (Fig. 4C) with dose-dependent inhibitory responses, thus resembling what has been observed in the *Xenopus* oocyte recording system (Fig. 1). Additionally, the responses to cyclohexanone were modulated by both eucalyptol (Fig. 4D) and fenchone (Fig. 4E) in a dose-dependent manner thus implying that ORN-A in SST-2 expresses CquiOR32.

**Fig. 4.**
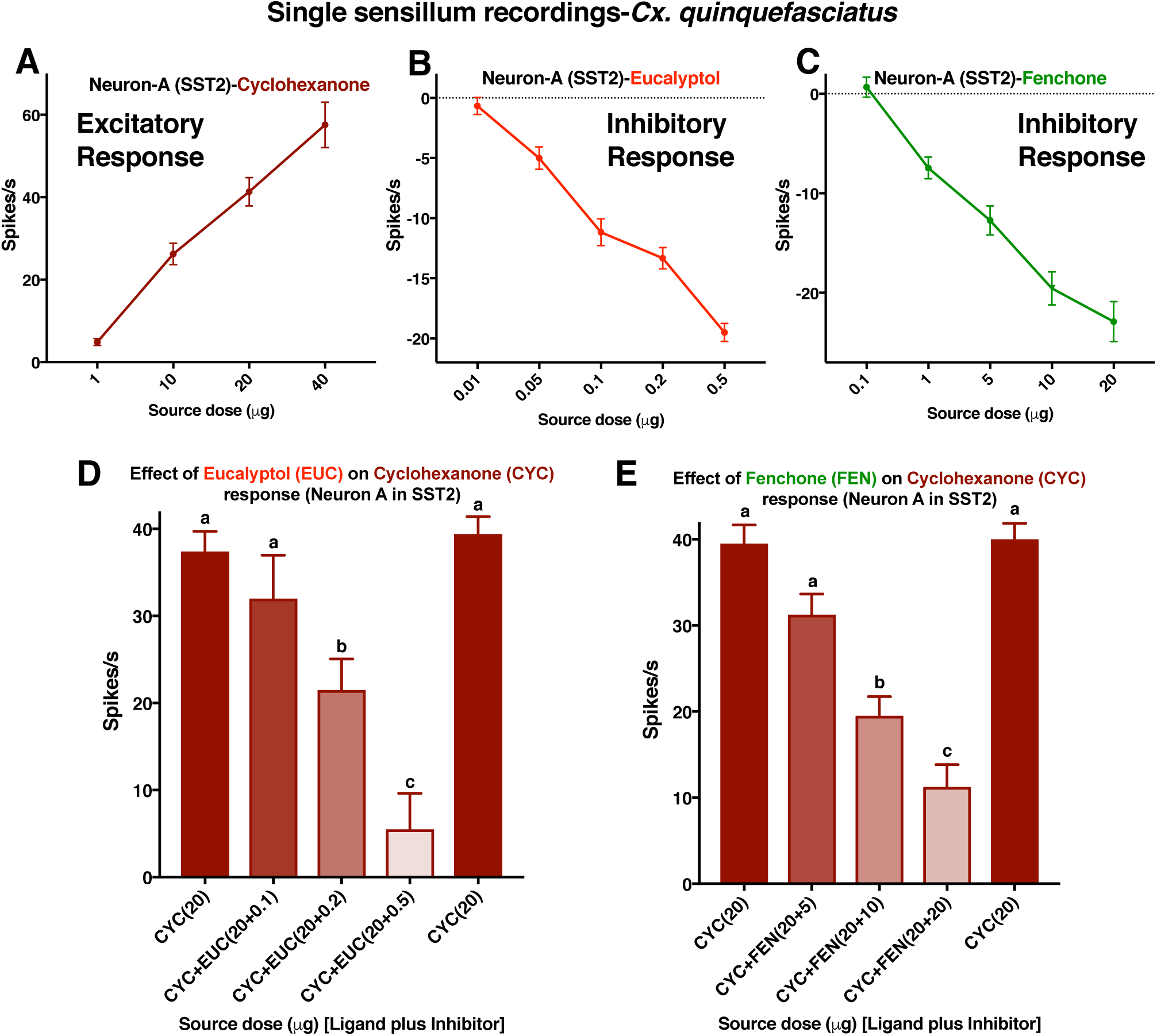
Single sensillum recordings from short sharp-tipped-2 (SST-2) sensilla on *Cx. quinquefasciatus* antennae. (A-C) Dose-dependent curves for excitatory and inhibitory compounds. (D, E) Effects of eucalyptol (EUC) and fenchone (FEN) on the excitatory responses elicited by cyclohexanone (CYC). Stimuli were delivered in sequence as displayed from left to right. Firing rates observed during 500 ms post-stimulus period were subtracted from spontaneous activities observed in the 500 ms pre-stimulus period and the outcome was multiplied by 2 to obtain the number of spikes per second. Error bars represent SEM. n = 6-12. Bars with different letters are considered statistically different at the 0.05 level, according to Tukey’s test.

### CquiOR32 variant with stronger inhibitory currents

We sequenced 30 CquiOR32 clones, of which 12 were identical and were then considered to be the wild type (WT). Their amino acid sequences showed 99.7% identity to CquiOR32 in VectorBase (CPIJ004156), the only difference being residue 249 (Ile in VectorBase vs. Val in our *Cx. quinquefasciatus* colony, GenBank, KM229539). Nine clones were pseudogenes and, therefore, were not tested.

Six clones differed from the WT in single amino acid residues. When tested in the *Xenopus* oocyte recording system, CquiOR32H161L (MG593061), CquiOR32Y350C (MG593062), and CquiOR32S127P (MG593064) behaved like WT, whereas CquiOR32W284R (MG593065) and CquiOR32M288T (MG593060) gave weak and very weak responses, respectively, and CquiOR32L299P (MG593066) showed moderate inward, but no inhibitory currents.

Two variants, differed in 2 amino acid residues, one, CquiOR32D62A,D119V (MG593067), behaving as WT and the other, CquiOR32D62A,R233L (MG593063), giving no response. One variant, CquiOR32D250_I251insIVELVD (MG593059) generated robust reverse currents and weak inward currents when challenged with excitatory and inhibitory compounds at the same dose. In addition to the insert (IVELVD), this variant differed from the WT in 2 amino acid residues, ie, F141L,V148A. Deletion of the insert generates the mutant CquiOR32I251_D256del, which generated a response profile indistinguishable from that of CquiOR32D250_I251insIVELVD.

Because CquiOR32I251_D256del differed from WT in 2 residues (F141L,V148A), we generated and tested 2 point mutations. Whereas CquiOR32I251_D256delL141F (only residue-141 rescued) showed inward and inhibitory current or relatively the same intensity, CquiOR32I251_D256delA148V (only residue-148 rescued) gave relatively stronger inward than inhibitory currents thereby suggesting that both residues are important for inhibitory currents (fig. S6).

### CquiOR32 orthologue in *Aedes aegypti* and intrareceptor inhibition

CquiOR32 has an orthologue in the genome of the yellow fever mosquito, AaegOR71 (AAEL017564), with 55.5% identity. We sequenced 20 clones and obtained 19 AaegOR71 sequences. Five clones showed sequences identical to the sequence in VectorBase and were, therefore, considered the WT. We expressed AaegOR71-WT in the *Xenopus* oocyte recording system (along with AaegOrco) and challenged the oocytes with compounds that elicited inward and inhibitory currents in CquiOR32. Cyclohexanone elicited inward currents, but the compounds generating the largest inward currents were 4,5-dimethylthiazole (DMT) and 2-methyl-2-thiazoline (2MT) (fig. S7); no response was observed with methyl salicylate. Although eucalyptol and fenchone did not elicit measurable inhibitory currents, these 2 compounds reduced AaegOR71 responses to cyclohexanone, DMT, and 2MT (fig. S7).

Four clones (AaegOR71-V2, GenBank MG593069) differed from WT in 7 amino acid residues and 3 clones (AaegOR71-V1, MG593068) differed from WT in 11 amino acid residues. They both showed weak responses to odorants when tested in the *Xenopus* oocyte recording system. The other 7 clones differed from the WT in 6-12 amino acid residues. AaegOR71-V5 (MG593071), AaegOR71-V14 (MG593074), and AaegOR71-V15 (MG593075) differed in 12, 9, and 11 amino acid residues, respectively, and none of them responded to odorants. AaegOR71-V8 (MG593072) differed in 10 amino acid residues and showed very weak response only to the Orco agonist VUAA-1. AaegOR71-V4 (MG593070), AaegOR71-V9 (MG593073), AaegOR71-V17 (MG593076) differed in 8, 10, and 6 amino acid residues, but gave weak to moderate responses to odorants. None of these variants elicited detectable inhibitory currents when challenged with eucalyptol or fenchone.

### Peripheral inhibition in the yellow fever mosquito antennae

Next, we tested whether intrareceptor inhibition might be manifested in vivo in the antennae of the yellow fever mosquito. SSR showed that cyclohexanone, 2-methyl-2-thiazoline, and 2,4-dimethylthiazole elicited dose-dependent excitatory responses in neuron-A in SST-2, whereas eucalyptol and fenchone showed inhibitory responses (fig. S8). When costimulated with 2-methyl-2-thiazoline and eucalyptol, the response to the odorant decreased markedly (fig. S9). We then analyzed the effect of inhibitors on the responses to the 3 odorants that caused excitatory responses. Both eucalyptol and fenchone inhibited the responses of ORN-A in SST-2 to cyclohexanone (Fig. 5A,B), 2-methyl-2-thiazoline (Fig. 5C, D), and 4,5-dimethylthiazole (Fig. 5E,F) in a dose-dependent manner. Taken together, these findings suggest that both ORN-A houses AaegOR71 and, more importantly, intrareceptor inhibition occurs in vivo.

**Fig. 5.**
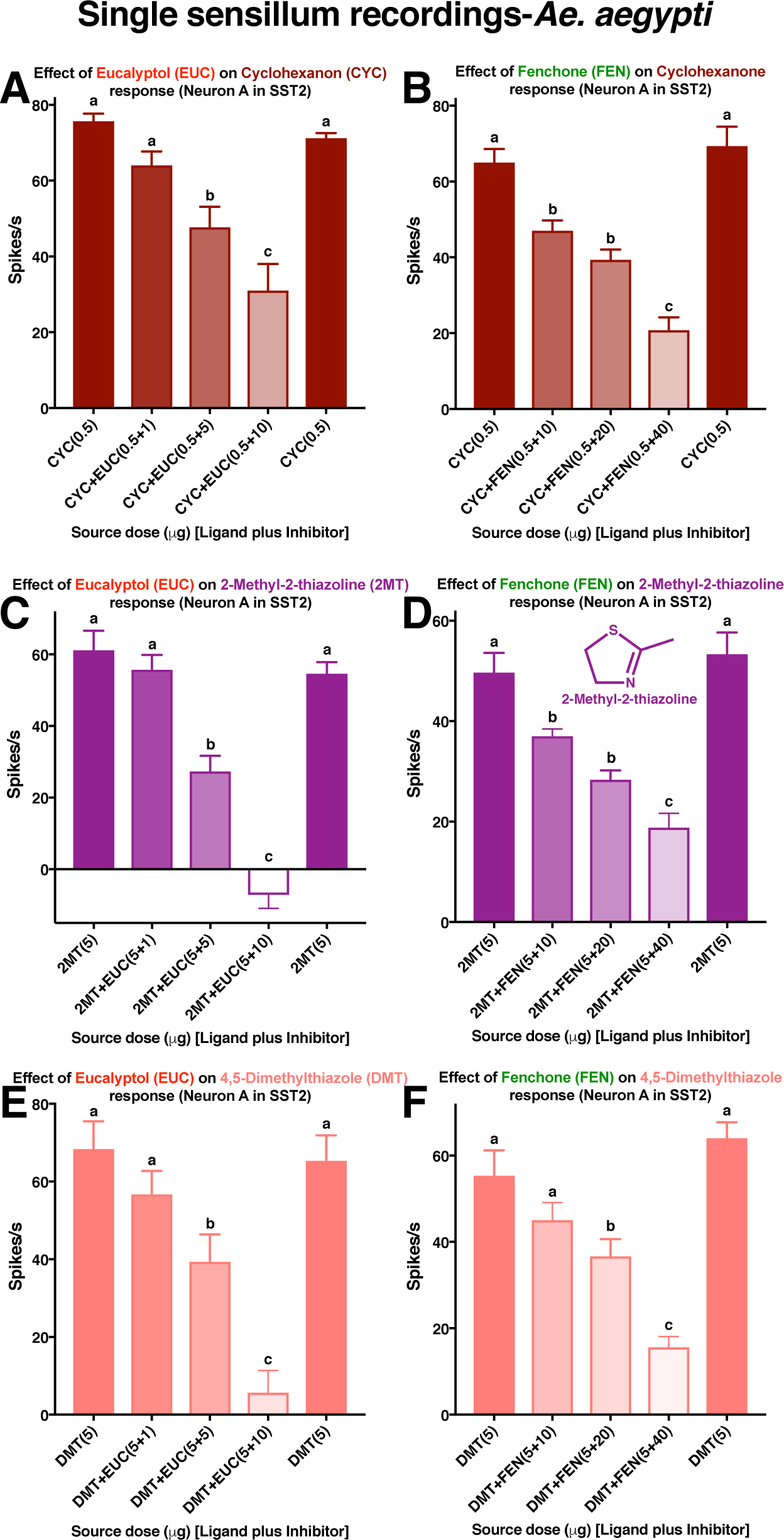
Quantification of responses from neuron-A in the short sharp-tipped-2 (SST-2) sensilla on *Ae. aegypti* antennae. The sensilla were challenges with mixtures of (A & B) cyclohexanone, (C &D) 2-methyl-2-thiazoline, or (E & F) 4,5-dimethylthiazole in combination with either eucalyptol (A, C & E) or fenchone (B, D & F). Stimuli were delivered in sequence as displayed from left to right. Firing rates observed during 500 ms post-stimulus period were subtracted from spontaneous activities observed in the 500 ms pre-stimulus period and the outcome was multiplied by 2 to obtain the number of spikes per second. Error bars represent SEM. n = 6.

### Intrareceptor inhibition manifested in mosquito behavior

Evidence in the literature suggests that eucalyptol is an oviposition deterrent *(14)*. We then tested in our surface landing and feeding assay (*32*) whether this inhibitory compound or other odorants would repel blood-seeking *Cx. quinquefasciatus* mosquitoes. At 0.1% eucalyptol, 47.57±3.15% mosquitoes responded to the treatment side of the arena, whereas 52.43±3.15% responded to the control (n = 10, P = 0.3276; eucalyptol, 7.5±0.4 mosquitoes in treatment and 8.5±0.8 mosquitoes in the control side). With a higher dose (1%), 44.3±5.7% and 55.7±5.7% mosquitoes responded to treatment and control, respectively (n = 10, P = 0.3330; treatment, 8.1±1.1; control 10.1±1.0 mosquitoes), thus showing that eucalyptol per se is not a potent spatial repellent. Likewise, cyclohexanone did not show repellence activity at 0.1% (47.7±4.2% treatment vs. 52.3±4.2% control, n = 4, P = 0.7027; 7±1.1 and 7.5±0.5 in treatment and control, respectively) or 1% (52.7±2.7% treatment vs. 47.3±2.7% control, n = 4, P = 0.4950; 8.25±0.9 and 7.75±1.4 in treatment and control, respectively).

Contrary to eucalyptol, methyl salicylate showed repellence activity. At a dose of 0.1%, the number of mosquitoes responding to the treatment side of the arena was significantly lower than the number responding to control (39.9±4.0% treatment vs. 60.1±4.0% control, n = 10, P = 0.0303; 8±0.8 and 12.3±1.1 in treatment and control, respectively). At a higher dose (1%), the difference was highly significant (15.3±3.7% treatment vs. 84.5±3.7% control, n = 10, P < 0.0001; 1.9±0.8 and 8.3±0.9 in treatment and control, respectively). Methyl salicylate (boiling point, 222 ^°^C) has a higher vapor pressure than DEET (boiling point, 285 ^°^C) and yet provided lower, albeit not significantly different, protection against mosquito bites than DEET did (Fig. 6). Therefore, it is unlikely a candidate for replacement of DEET in commercial applications.

**Fig. 6.**
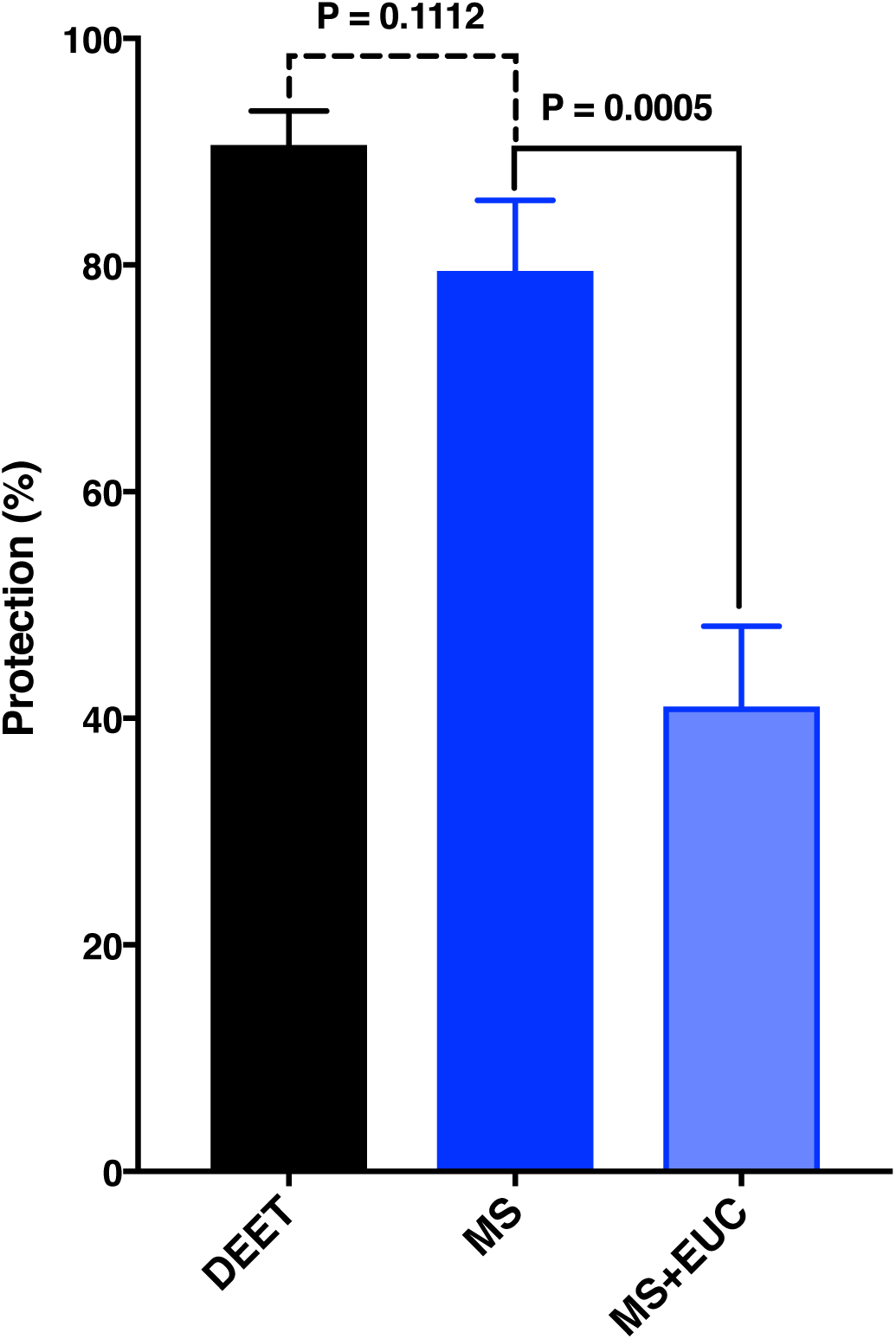
Effect of eucalyptol on methyl salicylate-elicited repellency. Although providing lower protection than DEET, methyl salicylate at the same dose (1%) repelled blood-seeking *Culex* mosquitoes. MS-elicited repellency was significantly reduced when tested in a mixture of 1% MS and 1% eucalyptol (EUC). Error bars represent SEM. n = 10.

Because eucalyptol per se is not a repellent, we then tested the effect of eucalyptol on repellence elicited by methyl salicylate. A mixture of 1% methyl salicylate and 1% eucalyptol had significantly lower protection than methyl salicylate alone (Fig. 6, Video 1). These behavioral measurements suggest that eucalyptol modulates the mosquito olfactory response to methyl salicylate, thus reducing the behavioral response. In short, the lower repellency observed with a mixture of methyl salicylate plus eucalyptol is equivalent to the repellency observed with a lower dose of methyl salicylate alone.

## CONCLUSIONS

We have serendipitously observed that ion channels formed by a mosquito odorant receptor along with its obligatory coreceptor Orco in the *Xenopus* oocyte recording system generate inward and inhibitory currents, which are likely correlated with the excitatory and inhibitory responses recorded by extracellular recordings (SSR) from antennal neurons. These inhibitory currents may lead to intrareceptor inhibition in vivo, which ultimately affects mosquito behavior.

## Acknowledgments

We thank Professor Karl-Ernst Kaissling (Max Planck Institute, Swiessen, Germany) for his comments on a draft version of the manuscript and former laboratory member, Mario S. Izidoro Júnior, for cloning one of the AaegOR71 variants. The National Institute of Allergy and Infectious Diseases (NIAID) of the National Institutes of Health under award numbers R01AI095514 and R21AI12893 supported research reported in this publication. The content is solely the responsibility of the authors and does not necessarily represent the official views of the NIH. F.Z. was supported by the Chinese Scholarship Council.

## Author contributions

P.X., Y.-M.C., Z.C., F.Z, and K.T. performed research; T.-Y.C., A.J.C., and N.L. contributed material and analytical tools; W.S.L. designed research, interpreted the data, and wrote the paper; all authors analyzed the data.

The authors declare no conflict of interest.

All raw data reported in this paper are included in the Data File S1.

## Supplementary Materials

### MATERIALS AND METHODS

#### Insect preparations and behavioral studies

*Cx. quinquefasciatus* used in this study were from a laboratory colony originating from adult mosquitoes collected in Merced, CA in the 1950s (*1*) and kept at the Kearney Agricultural Research Center, University of California, Parlier, CA. Specifically, we used mosquitoes from the UC Davis colony, which was initiated about 6 years ago with mosquitoes from the Kearney colony. In Davis, mosquitoes were maintained at 27±1^°^C, 75±5% relative humidity, and under a photoperiod of 12:12 h. Two colonies of *Ae. aegypti* were used in this study, namely, the Orlando strain, kept at Auburn University, and a colony established in 2016 from eggs that were laid by females collected in BG-sentinel traps (Biogents, Regensburgs, Germany) in the City of Clovis, California. These colonies were maintained at 25±2^°^C, 75±5% relative humidity, and under a photoperiod of 12:12 h. Repellency was measured using the previously reported surface landing and feeding assay (*2*). Behavioral responses are expressed in protection rate, according to WHO and EPA recommendations: P%=(1-[T/C]) X 100, where T and C represent the number of mosquitoes in treatment and control sides of the arena.

#### OR cloning

Total RNA samples were extracted from 1 thousand 4-7 day-old *Culex* female antennae and 800 *Aedes* female antennae with TRIzol reagent (Invitrogen, Carlsbad, CA). Antennal cDNA was synthesized from 1 μg of antennal total RNA from each species using a SMARTer™ RACE cDNA amplification kit according to manufacturer’s instructions (Clontech, Mountain View, CA). To obtain full-length coding sequences of CquiOR32, PCRs were performed using gene-specific primers containing restriction endonuclease sites and Kozak motif (acc): CquiOR32 Fwd-XmaI (underlined) primer, 5’-TCCC<u>CCCGGG</u>GGGAaccATGTTCACCACTCAAAACAGTCACC-3’and Rev-XbaI (underlined) primer, 5’-GC<u>TCTAGA</u>GCTTATATCATAGTCAACGCTTCCTTCAGCA-3’. PCR products were purified by a QIAquick gel extraction kit (Qiagen) and then cloned into pGEM-T vector (Promega). Plasmids were extracted using a QIAprep spin mini prep kit (Qiagen) and sequenced (Davis Sequencing). To subclone CqOR32 into pGEMHE, pGEM-T-CquiOR32 was digested by XmaI and XbaI (BioLabs) before being subcloned. After transformation, plasmids were extracted using the QIAprep Spin Miniprep kit (Qiagen) and sequenced by ABI 3730 automated DNA sequencer at Davis Sequencing (Davis, CA) for confirmation. Likewise, AaedOR71 was cloned with a pair of cloning primers: AaeagOR71-Fwd: AGATCAATTCCCCGGGaccATGGAACTGTCCTACCATCGAAGTC AaegOR71 -Rev: TCAAGCTTGCTCTAGACTAAAGCCGATCCAGAACATCTTTG

#### Quantitative PCR, qPCR

For qRT-PCR, each type of tissue (antennae, maxillary palps, proboscis, and legs) from 300 blood-fed female mosquitoes (4–7 days old) was dissected and collected in TRIzol reagent (Invitrogen, Carlsbad, CA) on ice using a stereo microscope (Zeiss, Stemi DR 1663, Germany). Total RNA was extracted using TRIzol reagent. After RNA was quantified on NanoDrop Lite spectrometer (Thermo Fisher Scientific, Rockford, IL), cDNA was synthesized from 200 ng of equal amount RNA using iScript™ Reverse Transcription Supermix for RT-qPCR according to the manufacturer’s instructions (Bio-Rad, Hercules, CA). Real-time quantitative PCR (qPCR) was carried out by using a CFX96 Touch™ Real-Time PCR Detection System (Bio-Rad) and SsoAdvanced SYBR Green Supermix (Bio-Rad). *CquiRPS7* gene was used as the reference. The following primers, designed by Primer 3 program (http://frodo.wi.mit.edu/), were used: CquiOR32-Fw: 5’-GCGATTTTTGCTTCGAAAAG-3’

CquiOR32-Rv: 5’-GTGCGTCCAATACCGAAAGT-3’.

qPCR was performed with 3 biological replicates, and each of them was replicated 3 times (3 technical replicates per biological replicate); data were analyzed using the 2–ΔΔCT method.

#### Electrophysiology

Tested flies were selected from F1 progeny for *D. melanogaster*, which carried both UAS-CquiOR36 and DmelOrco-Gal4 promoter (*3*). The EAG apparatus (Syntech Ltd., Hilversum, The Netherlands) was linked to a computer with an EAG2000 data acquisition interface. Recording and indifferent electrodes were made of Ag/AgCl wires enclosed in drawn glass capillary needles, which were filled with 1 M potassium chloride in 1% polyvinylpyrrolidone. The reference electrode was inserted in the eye of an immobilized insect and the recording electrode was placed on the third segment of a fruit fly antenna by using a micromanipulator MP-12 (Syntech). Compounds used as stimuli were freshly dissolved in paraffin oil and loaded on a filter paper strip (1 cm^2^), which were placed into Pasteur pipettes as cartridges. The preparation was bathed in a high-humidity air stream flowing from a Stimulus Controller CS-55 (Syntech) at 160 mL/min to which compensatory flow or stimulus pulse (125 mL/s, 300 ms) was added. For dual delivery, the stimulus flow was split and passed through 2 cartridges, each one having a filter paper strip laden with one of the tested compounds. The outlets of these cartridges merged in the continuous flow and placed 1 cm away from the antennal preparation. Signal from the antenna induced by stimulus or control puff was recorded for 10 s.

For SSR, 4- to 5-day-old female mosquitoes were used after being anesthetized on ice and fixed with a 200-μL pipette tip (*6*). Mosquitoes were fixed with dental wax and using a cover slip (22 X 22 mm) with double-sided tape. The reference tungsten electrode was inserted into one eye of a test mosquito, whereas the recording electrode was inserted into the shaft of a test sensillum under a microscope (Leica Z6 Apo) by using a micromanipulator (Leica, Cat #: 115378). Chemical compounds used as stimuli were freshly prepared with dimethyl sulfoxide (DMSO) at desired concentrations and 1 μL of each chemical solution was dispersed onto a piece of filter paper (3 × 45 mm), which was inserted into a glass Pasteur pipette to create a stimulus cartridge. The preparation was bathed in a high-humidity air stream flowing from a Stimulus Controller CS-55 (Syntech) at 20 mL/s to which compensatory flow or stimulus pulse (0.5 L/min, 500 ms) was added. The signal acquired by a preamplifier (Universal AC/DC Probe Gain 10X, Syntech) was digitized by using an IDAC 4 (Syntech). Action potential evoked by stimulus or control puff was recorded for 10 s, starting 1 s before the stimulation. Action potentials were counted off-line over a 500-ms period before and after the stimulation (*4*). Specifically, firing rates observed during 500 ms post-stimulus were subtracted from spontaneous activities observed in the 500 ms pre-stimulus period and the outcome was multiplied by 2 to obtain the number of spikes per second (*4*).

The 2-electrode voltage-clamp technique (TEVC) was performed as previously described (*5-9*). Briefly, the capped cRNAs were synthesized using pGEMHE vectors and mMESSAGE mMACHINE T7 Kit (Ambion). Purified OR cRNAs were resuspended in nuclease-free water at 200 ng/mL and 9.2 nl aliquots were microinjected with the same amount of CquiOrco cRNA into *Xenopus laevis* oocytes in stage V or VI (purchased from EcoCyte Bioscience, Austin, TX). Then, the oocytes were kept at 18^o^C for 3–7 days in modified Barth’s solution (NaCl 88 mM, KCl 1 mM, NaHCO_3_ 2.4 mM, MgSO_4_ 0.82 mM, Ca(NO_3_)_2_ 0.33 mM, CaCl_2_ 0.41 mM, HEPES 10 mM, pH 7.4) supplemented with 10 mg/mL of gentamycin, 10 mg/mL of streptomycin. Odorant-induced currents at holding potential of −80 mV were collected from oocytes bathed in perfusion Ringer (NaCl 96 mM, KCl 2 mM, CaCl_2_ 1.8 mM, MgCl_2_ 1 mM, HEPES 5 mM. pH 7.6) flowing at 3.2 ml/min. Stimulus were injected (100 μl in 2 s) at 1 cm upstream of the flow. After each stimulus, oocytes were thoroughly washed until a steady baseline was recovered. Source doses rather than the actual doses reaching oocytes are reported. Currents were amplified with an OC-725C amplifier (Warner Instruments, Hamden, CT), low-pass filtered at 50 Hz and digitized at 1 kHz. Data acquisition and analysis were carried out with Digidata 1440A and pCLAMP 10 software (Molecular Devices, LLC, Sunnyvale, CA). The panel of odorants included the following compounds: 1-butanol, 1-pentanol, 1-hexanol, 1-heptanol, 1-octanol, 1-nonanol, 1-dodecanol, 2,3-butanediol, 2-butoxyethanol, 3-methyl-1-butanol, 2-hexen-1-ol, 3-hexen-1-ol, 1-hexen-3-ol, 1-hepten-3-ol, 3-octanol, 1-octen-3-ol, 1-octyn-3-ol, 3-octyn-1-ol, 2-octanol, 2-butanol, 2-nonen-1-ol, 2-pentanol, 4-methylcyclohexanol, 1-hexadecanol, 3-pentanol, 3-methyl-2-butanol, 3-methyl-2-buten-1-ol, 2-methyl-3-buten-2-ol, p-menthane-3,8-diol (PMD), propanal, butanal, penatanal, isovaleraldehyde, hexanal, (E)-2-methyl-2-butenal, heptanal, octanal, nonanal, decanal, undecanal, 1-dodecanal, (E)-2-hexenal, (Z)-8-undecenal, (E)-2-heptenal, (E)-2-nonenal, phenylacetaldehyde, 2,4-hexadienal, furfural, benzaldehyde, a-hexylcinnamaldehyde, methyl acetate, ethyl acetate, propyl acetate, butyl acetate, pentyl acetate, hexyl acetate, heptyl acetate, octyl acetate, nonyl acetate, decyl acetate, methyl, propionate, ethyl propionate, methyl butyrate, ethyl butanoate, methyl hexanoate, ethyl hexanoate, ethyl 3-hydroxyhexanoate, ethyl 3-hydroxybutanoate, ethyl linoleate, phenyl propanoate, phenethyl propionate, ethyl 2-(E)-4-(Z)-decadienoate, (E)-2-hexenyl acetate, (Z)-3-hexenyl acetate, (E)-2-hexenyl acetate, ethyl lactate, phenyl isobutyrate, eugenyl acetate, methyl salicylate, ethyl stearate, methyl myristate, isopropyl myristate, palmitic acid methyl ester, 1-octen-3-yl acetate, isopentyl acetate, ethyl phenylacetate, geranyl acetate, octadecyl acetate, acetylacetone, 2-butanone, 2-heptanone, geranylacetone, 6-methyl-5-hepten-2-one, 5-methyl-2-hexanone, 2,3-butanedione, 3-hydroxy-2-butanone, 2-pentanone, 2-hexanone, 2-octanone, 2-undecanone, 2-tridecanone, 2-nonanone, 1-octen-3-one, cyclohexanone, acetophenone, γ-valerolactone, γ-hexalactone, γ-octalactone, γ-decalactone, γ-dodecalactone, p-coumaric acid, isovaleric acid, dodecanoic acid, (±)-lactic acid, ethanoic acid, propanoic acid, butanoic acid, isobutyric acid, 2-oxobutyric acid, pentanoic acid, 2-oxovaleric acid, myristic acid, palmitoleic acid, oleic acid, hexanoic acid, (E)-2-hexenoic acid, 5-hexanoic acid, (E)-3-hexenoic acid, heptanoic acid, octanoic acid, nonanoic acid, decanoic acid, n-tridecanoic acid, linoleic acid, ammonia, trimethylamine, triethylamine, propylamine, butylamine, pentylamine, hexylamine, heptylamine, octylamine, 1,4-diaminobutane, cadaverine, 1,5-diaminopentane, phenol, 2-methylphenol, 3-methylphenol, 4-methylphenol, 4-ethylphenol, 3,5-dimethylphenol, 2,3-dimethylphenol, 2,4-dimethylphenol, 2,5-dimethylphenol, 2,6-dimethylphenol, 3,4-dimethylphenol, guaiacol, 2-methoxy-4-propylphenol, 2-phenoxyethanol, 1,2-dimethoxybenzene, benzyl alcohol, 2-phenylethanol, 1-phenylethanol, phenylether, (S)-(-)-perillaldehyde, fenchone, thujone, camphor, α-terpinene, γ-terpinene, (-)-menthone, menthyl acetate, limonene, linalyl acetate, α-humulene, linalool oxide, geraniol, nerol, thymol, (±)-linalool, eucalyptol, citral, eugenol, α-pinene, ocimene, (±)-citronellal, α-phellandrene, nerolidol, jasmine, menthol, carvone, cymene, sabinene, terpinolene, ß-myrcene, (+)-δ-cadinene, (+)-limonene oxide, (*E,E*)-farnesol, (*E,E*)-farnesyl acetate, farnesene, α-methylcinnamaldehyde, cinnamyl alcohol, α-terpineol, citronellol, (*E*)-cinnamaldehyde, (-)-caryophyllene oxide, ß-caryophyphyllene, carvacrol, terpinen-4-ol, 7-hydroxcitronellal, N-(2-isopropylphenyl)-3-methylbenzamide, N-sec-butyl-2-phenyl-acetamide, N-(sec-butyl)-2-methylbenzamide, N-methylbenzamide, m-toluamide, (N,N)diethyl-3-methylbenzamide, pyridine, pyrrolidine, 2-pyrrolidinone, indole, 1-methylindole, 2-methylindole, 3-methylindole (skatole), 4-methylindole, 5-methylindole, 6-methylindole, 7-methylindole, isoprene, carbon disulfide, dimethyl trisulfide, 5-isobutyl-2,3-dimethylpyrazine, dibutyl phthalate, dimethyl phthalate, phenethyl formate, benzyl formate, 2-acetylthiophene, methyl disulfide, 2-ethyltoluol, 2-methyl-2-thiazoline, 2,4-thiazolinedione, 3-methylbenzamide, *N,N*-diethyl-3-methylbenzamide (DEET), ethyl *N*-acetyl-*N*-butyl-β-alaninate (IR3535), butan-2-yl 2-(2-hydroxyethyl)piperidine-1-carboxylate (picaridin), methyl anthranilate, methyl *(N,N)-*dimethylanthranilate, 4,5-dimethylthiazole, 4-dimethylamino-1-naphthaldehyde, and N-(4-ethylphenyl)-2-[(4-ethyl-5-pyridin-3-yl-1,2,4-triazol-3-yl)sulfanyl]acetamide (VUAA-1).

---

**Fig. S1.**
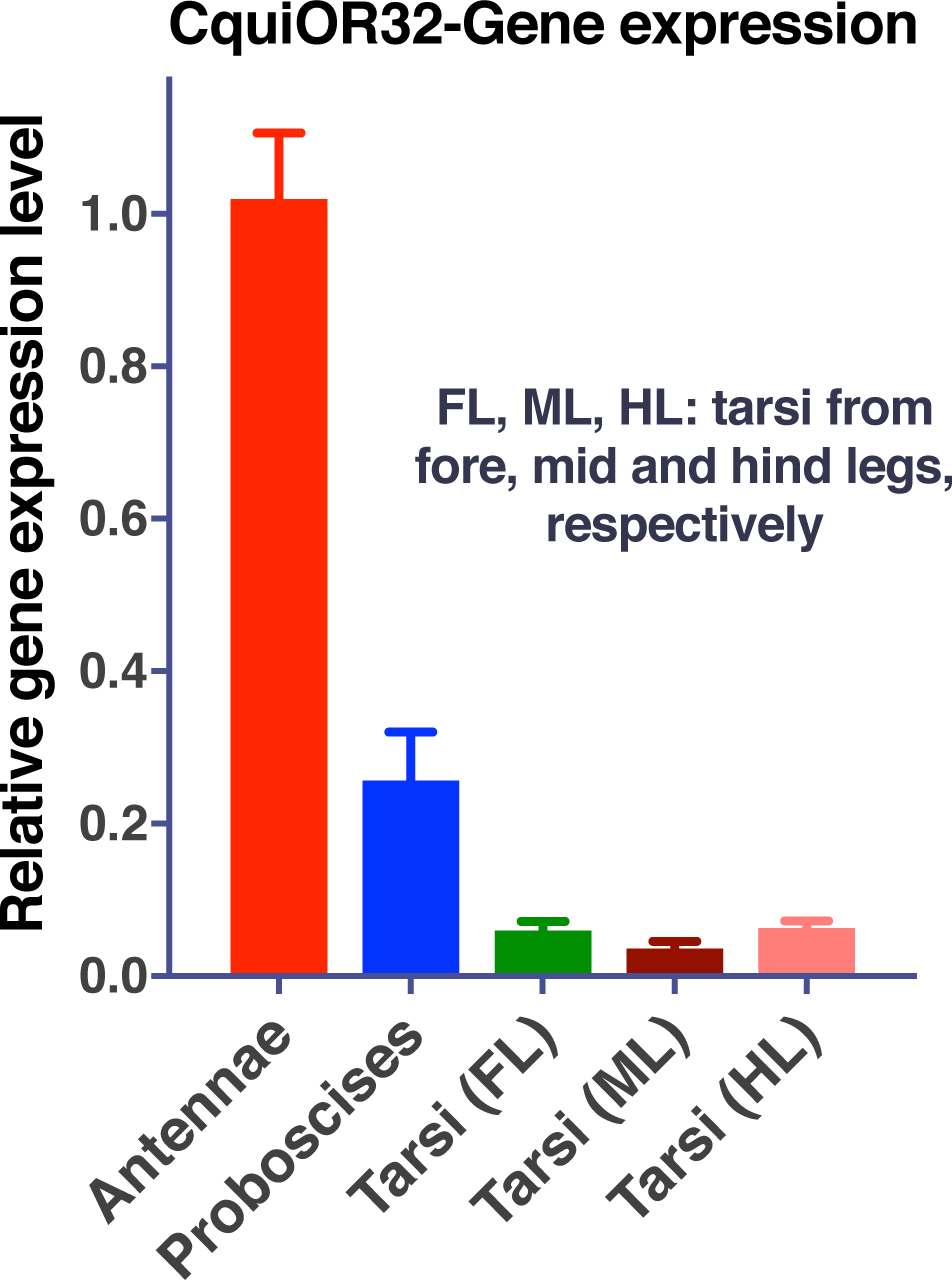
Quantitative PCR data for CquiOR32. This receptor is predominantly expressed in female antennae. Error bars represent SEM. n = 3 biological samples, each with 3 biological replicates.

**Fig. S2.**
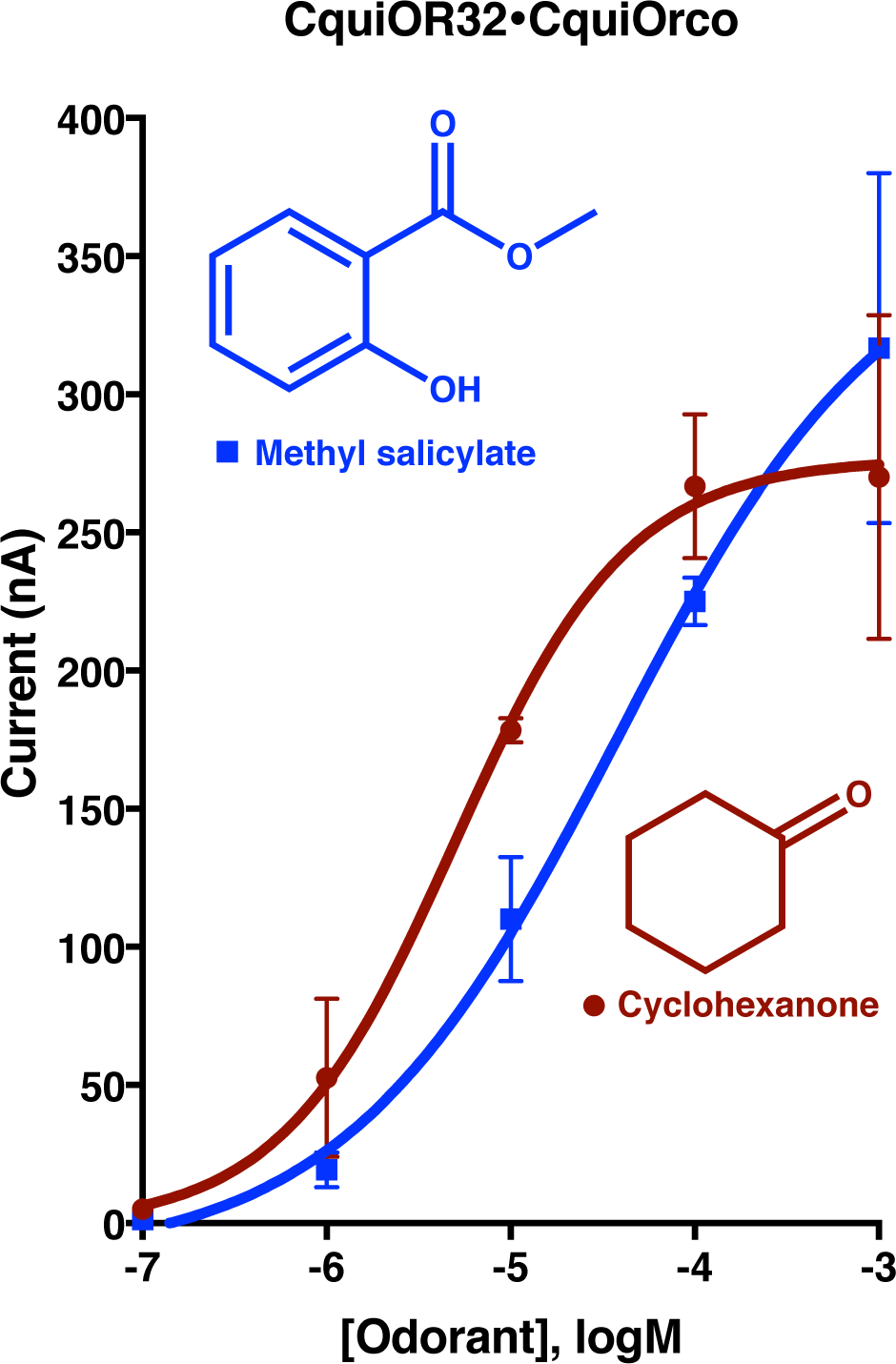
Concentration-response relationships. EC_50_ for methyl salicylate and cyclohexanone: 3.7×10^−5^M and 4.9×10^−6^M, respectively. N = 3 for each data point, each from a different oocyte of the same batch of eggs. Error bars represent SEM.

**Fig. S3.**
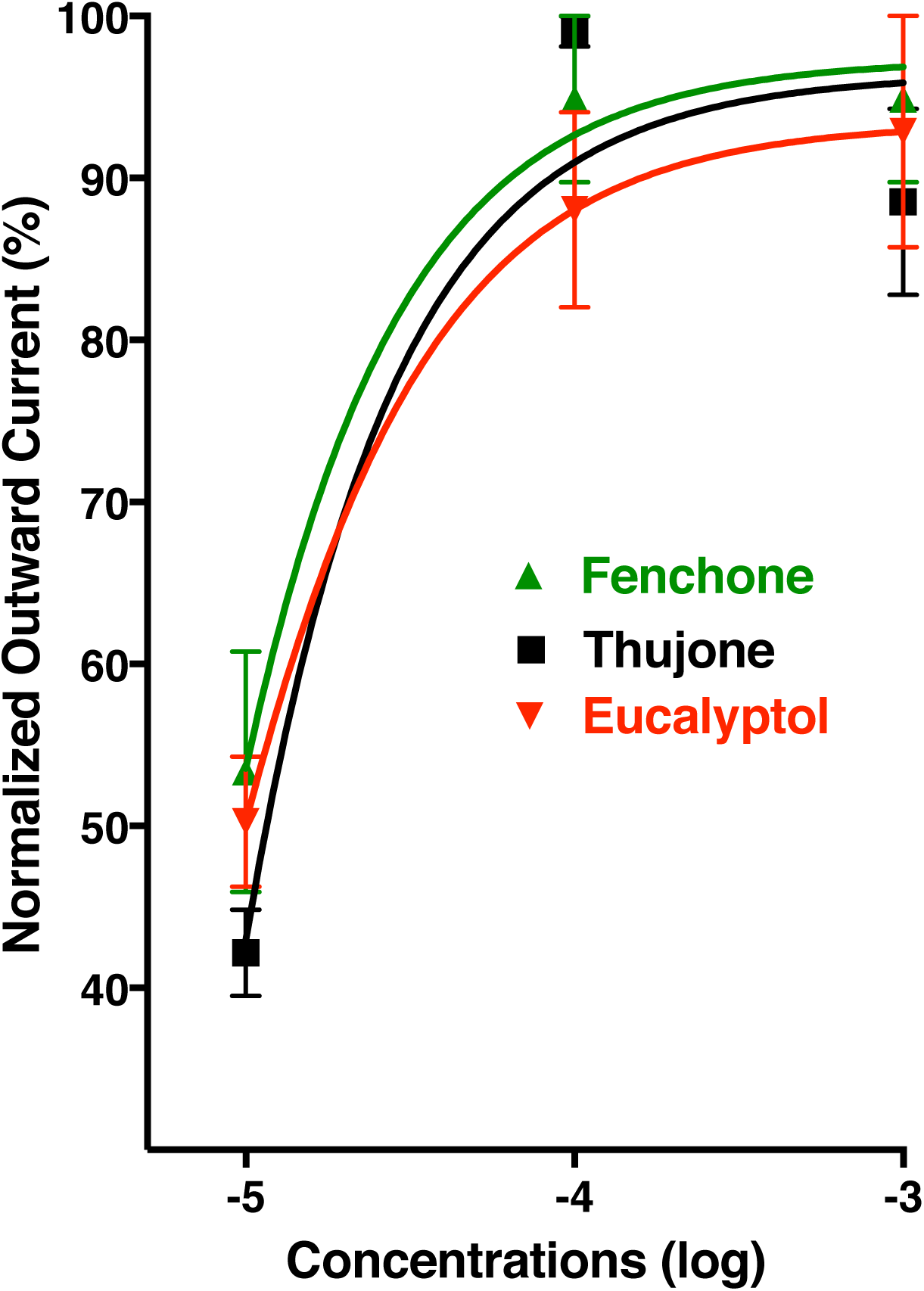
Dose-dependence curves for inhibitory compounds. Responses were normalized (n = 3 for each compound and dose). Error bars represent SEM.

**Fig. S4.**
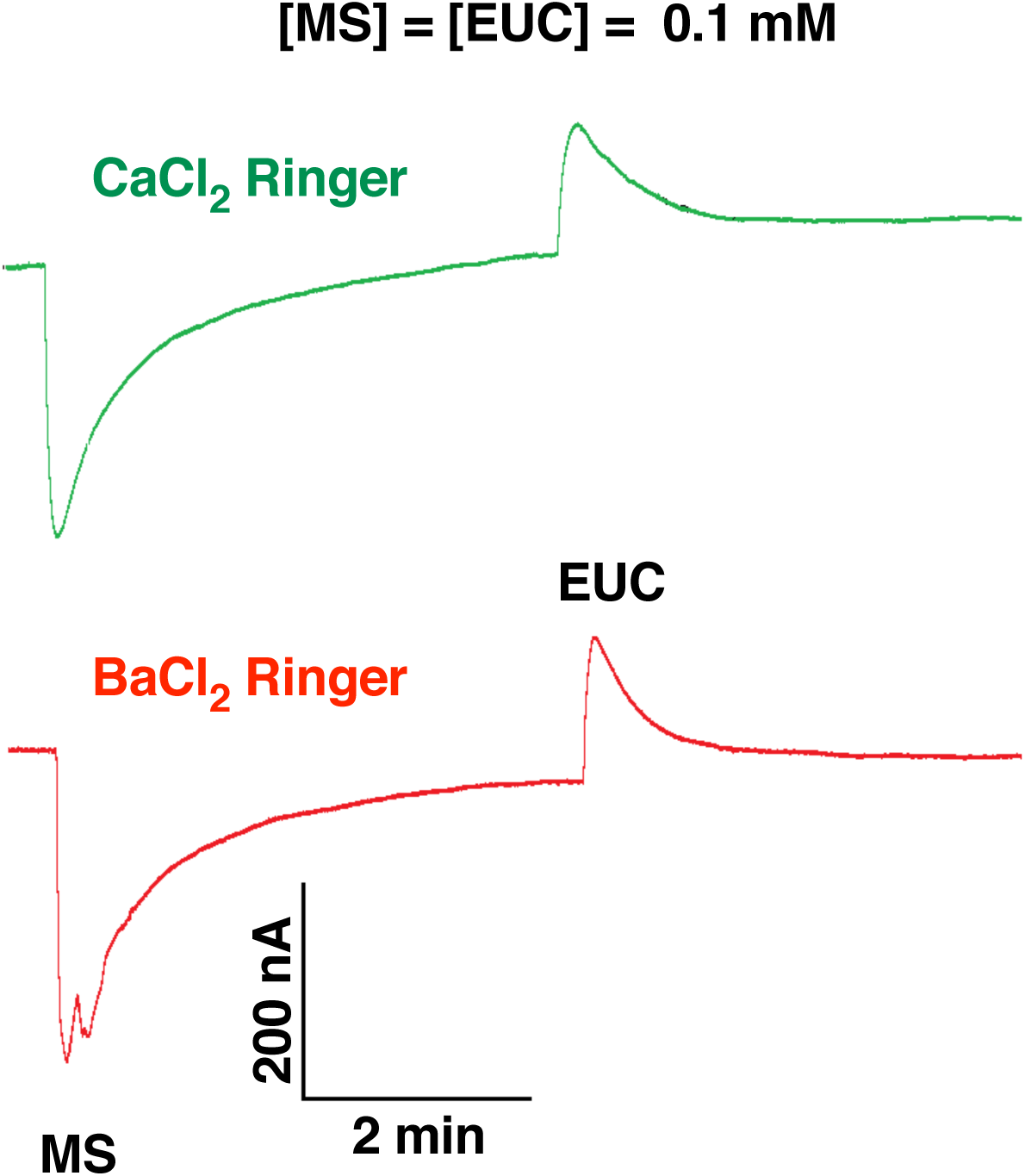
Currents recorded from CquiOR32-CquiOrco-expressing with regular buffer and a Ringer buffer having BaCl_2_ replacing CaCl_2_. Traces from the same oocyte bathed initially in regular buffer and then in a Ringer buffer with BaCl_2_ instead of CaCl_2_.

**Fig. S5.**
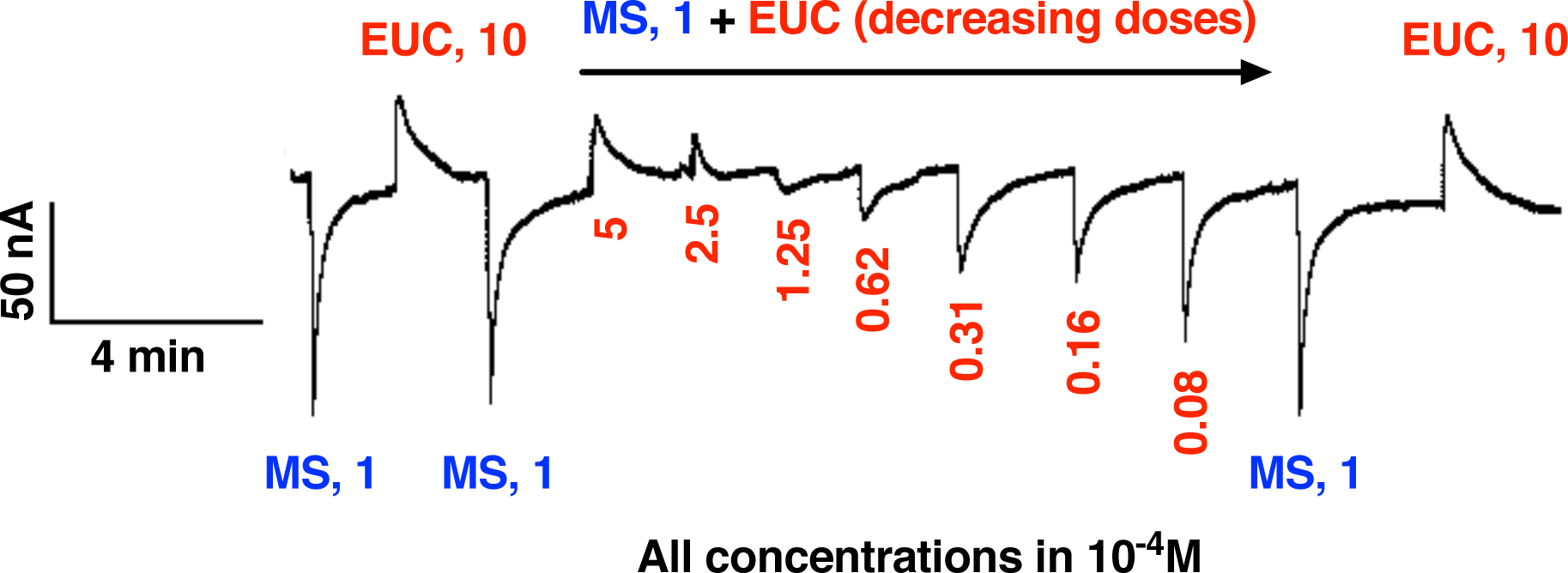
Eucalyptol (EUC)-elicited, dose-dependent inhibition of responses of CquiOR32-CquiOrco-expressing oocytes to methyl salicylate (MS). Continuous trace from the same oocyte.

**Fig. S6.**
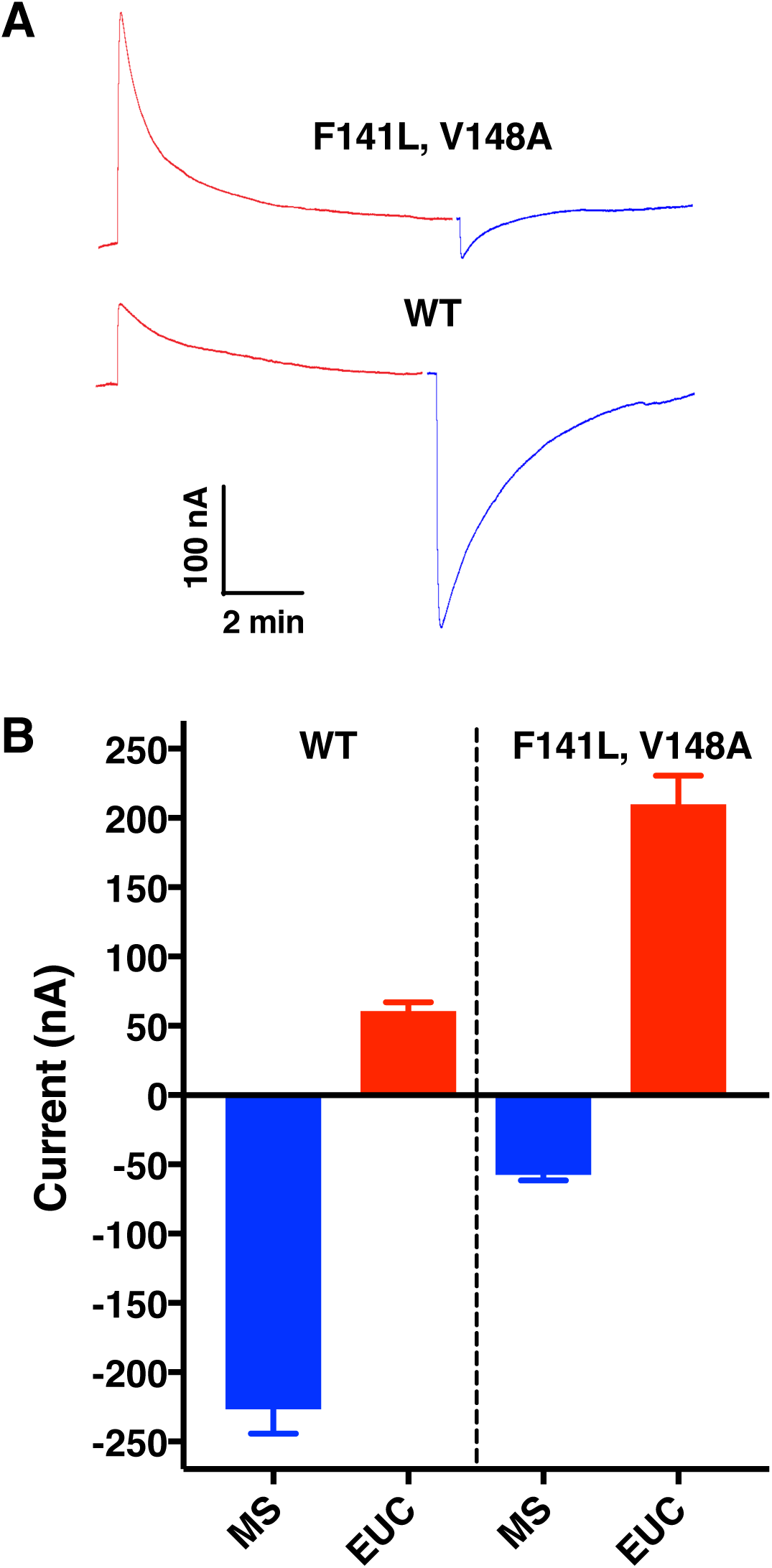
Currents elicited by CquiOR32-CquiOrco-expressing oocytes. (A) As opposed to WT, the channels formed with a natural variant (CquiOR32D250_I251insIVELVD) responded to eucalyptol with robust inhibitory current, whereas methyl salicylate generated smaller inward currents. (B) Quantification of responses. Error bars represent SEM. n = 3.

**Fig. S7.**
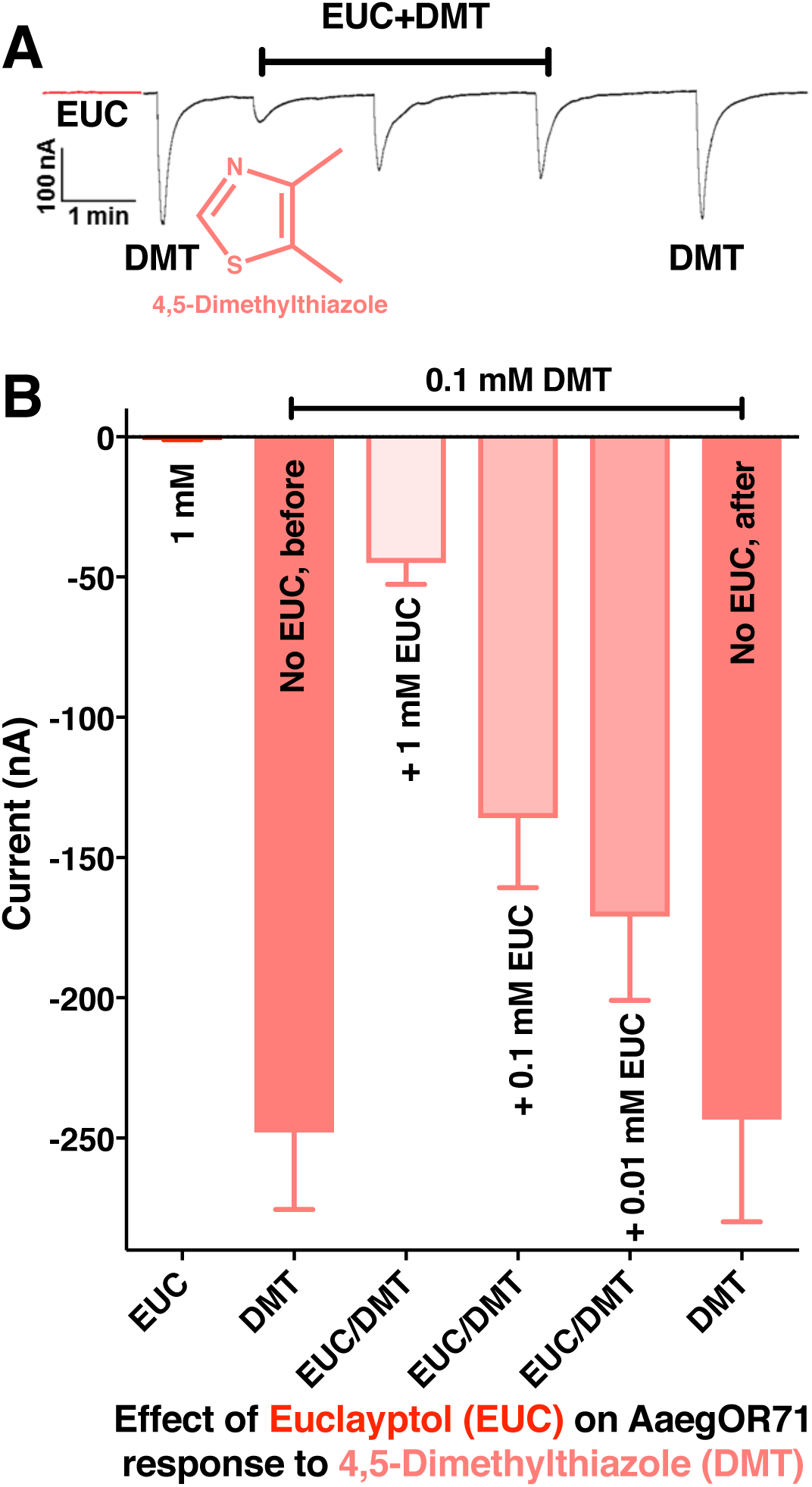
Eucalyptol inhibition of responses elicited by 4,5-dimethylthiazole (DMT) on oocytes coexpressing AaegOR71 and AaegOrco. Although eucalyptol (EUC) did not elicit detectable inhibitory currents, it caused a dose-dependence reduction in DMT-elicited responses. Error bars represent SEM. n = 5. Bars with different letters are considered statistically different at the 0.05 level, according to Tukey’s test.

**Fig. S8.**
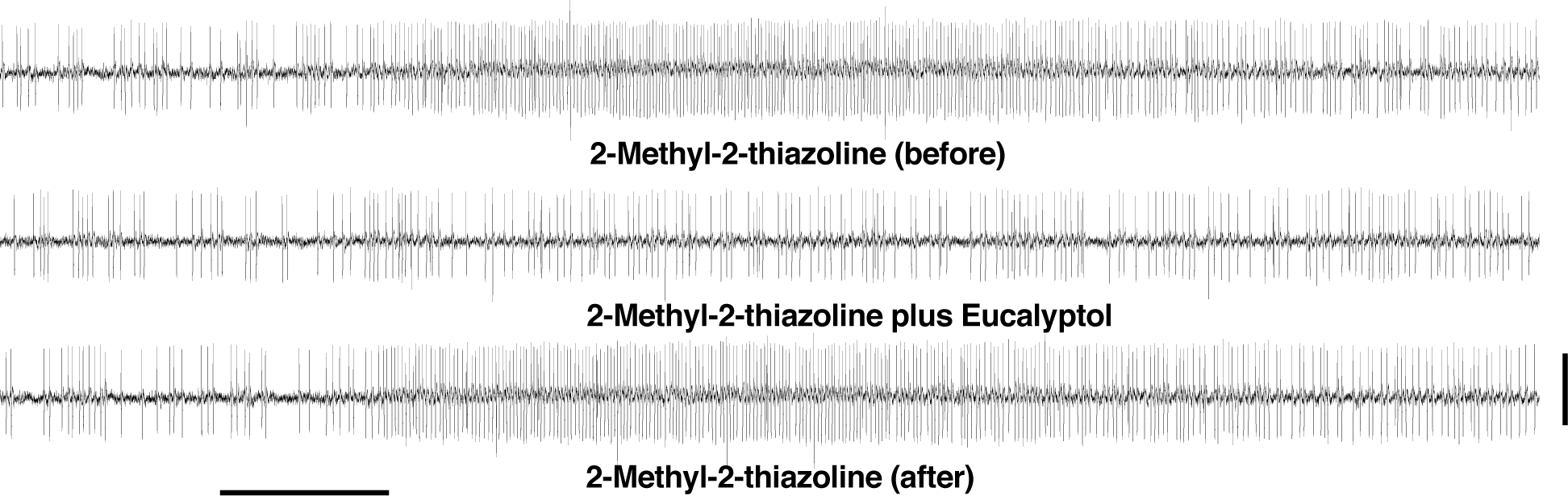
SSR traces from SST-2 sensilla on *Ae. aegypti* antennae. A bar beneath the traces indicates the duration of the stimulus (500 ms). Horizontal line after the bottom trace, 4 mV.

**Fig. S9.**
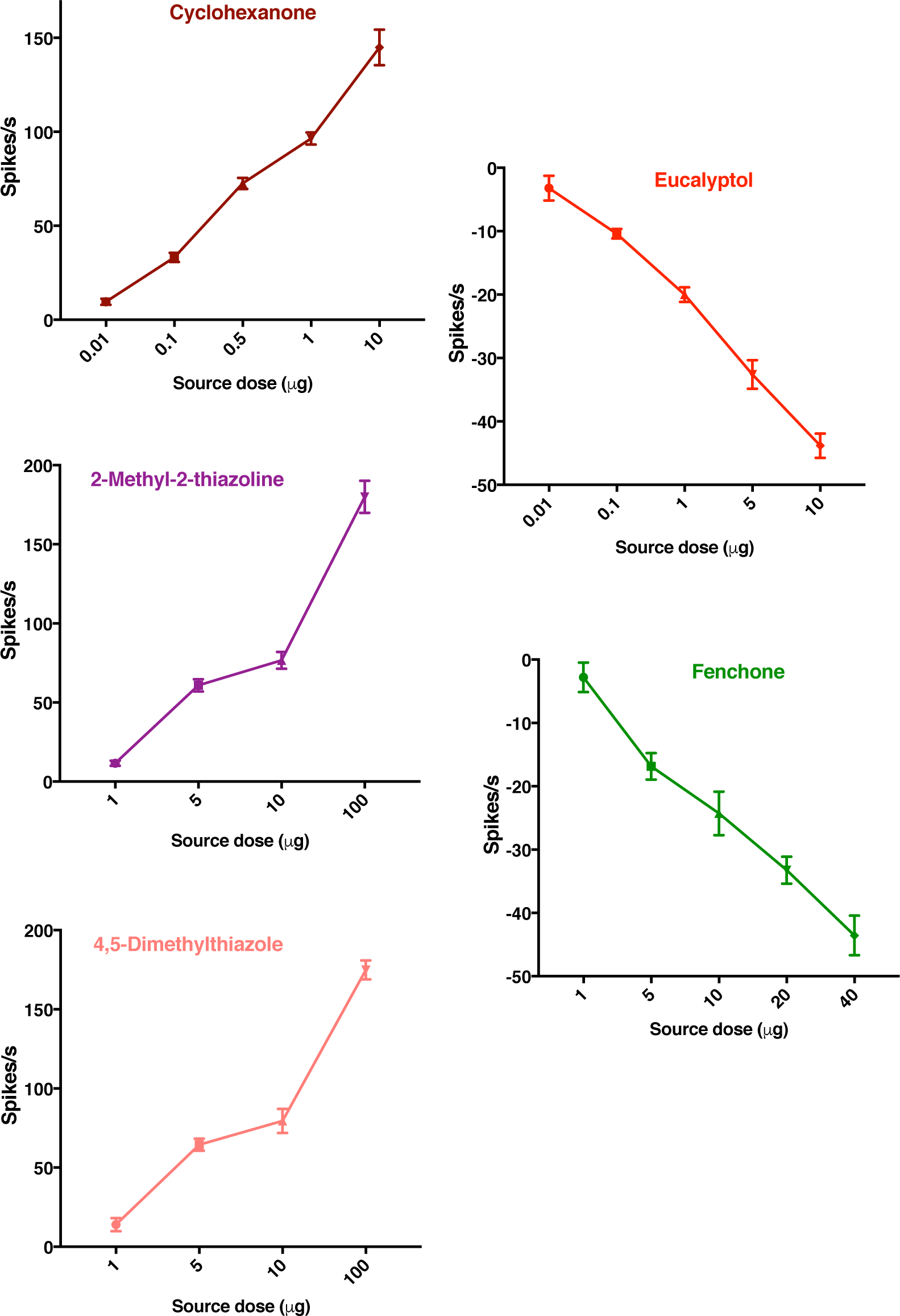
Dose-dependent curves recorded from *Ae. aegypti* antennae by SSR. Firing rates observed during 500 ms post-stimulus period were subtracted from spontaneous activities observed in the 500 ms pre-stimulus period and the outcome was multiplied by 2 to obtain the number of spikes per second. Error bars represent SEM. n = 5-20.

## REFERENCES AND NOTES

1. R. M. Joseph, J. R. Carlson, *Drosophila* chemoreceptors: A molecular interface between the chemical world and the brain. Trends Genet 31, 683–695 (2015).

2. S. R. Hill, B. S. Hansson, R. Ignell, Characterization of antennal trichoid sensilla from female southern house mosquito, *Culex quinquefasciatus* Say. Chem Senses 34, 231–252 (2009).

3. Z. Syed, W. S. Leal, Acute olfactory response of *Culex* mosquitoes to a human- and bird-derived attractant. Proc Natl Acad Sci U S A 106, 18803–18808 (2009).

4. F. Liu, L. Chen, A. G. Appel, N. Liu, Olfactory responses of the antennal trichoid sensilla to chemical repellents in the mosquito, *Culex quinquefasciatus*. J Insect Physiol 59, 1169–1177 (2013).

5. Z. Ye, F. Liu, N. Liu, Olfactory responses of southern house mosquito, *Culex quinquefasciatus*, to human odorants. Chem Senses 41, 441–447 (2016).

6. M. Ghaninia, R. Ignell, B. S. Hansson, Functional classification and central nervous projections of olfactory receptor neurons housed in antennal trichoid sensilla of female yellow fever mosquitoes, *Aedes aegypti*. Eur J Neurosci 26, 1611–1623 (2007).

7. K. E. Kaissling, Peripheral mechanisms of pheromone reception in moths. Chem Senses 21, 257–268 (1996).

8. A. A. Nikonov, W. S. Leal, Peripheral coding of sex pheromone and a behavioral antagonist in the Japanese beetle, *Popillia japonica*. J Chem Ecol 28, 1075–1089 (2002).

9. C. Y. Su, K. Menuz, J. Reisert, J. R. Carlson, Non-synaptic inhibition between grouped neurons in an olfactory circuit. Nature 492, 66–71 (2012).

10. G. M. Tauxe, D. MacWilliam, S. M. Boyle, T. Guda, A. Ray, Targeting a dual detector of skin and CO2 to modify mosquito host seeking. Cell 155, 1365–1379 (2013).

11. M. G. de Brito Sanchez, K. E. Kaissling, Inhibitory and excitatory effects of iodobenzene on the antennal benzoic acid receptor cells of the female silk moth *Bombyx mori* L. Chem Senses 30, 435–442 (2005).

12. W. V. van Naters, Inhibition among olfactory receptor neurons. Frontiers in Human Neuroscience 7, (2013).

13. P. Xu, Y. M. Choo, A. De La Rosa, W. S. Leal, Mosquito odorant receptor for DEET and methyl jasmonate. Proc Natl Acad Sci U S A 111, 16592–16597 (2014).

14. J. A. Klocke, M. V. Darlington, M. F. Balandrin, 1,8-Cineole (Eucalyptol), a mosquito feeding and ovipositional repellent from volatile oil of *Hemizonia fitchii* (Asteraceae). J Chem Ecol 13, 2131–2141 (1987).

15. D. Wicher et al., *Drosophila* odorant receptors are both ligand-gated and cyclic-nucleotide-activated cation channels. Nature 452, 1007–1011 (2008).

16. R. Smart et al., *Drosophila* odorant receptors are novel seven transmembrane domain proteins that can signal independently of heterotrimeric G proteins. Insect Biochem Mol Biol 38, 770–780 (2008).

17. K. Sato et al., Insect olfactory receptors are heteromeric ligand-gated ion channels. Nature 452, 1002–1006 (2008).

18. C. Hartzell, I. Putzier, J. Arreola, Calcium-activated chloride channels. Annu Rev Physiol 67, 719–758 (2005).

19. D. Wicher, Olfactory signaling in insects. Prog Mol Biol Transl Sci 130, 37–54 (2015).

20. D. W. McBride, Jr., S. D. Roper, Ca(2+)-dependent chloride conductance in *Necturus* taste cells. J Membr Biol 124, 85–93 (1991).

21. R. Taylor, S. Roper, Ca(2+)-dependent Cl-conductance in taste cells from *Necturus*. J Neurophysiol 72, 475–478 (1994).

22. R. A. Steinbrecht, Experimental morphology of insect olfaction: tracer studies, X-ray microanalysis, autoradiography, and immunocytochemistry with silkmoth antennae. Microsc Res Tech 22, 336–350 (1992).

23. K. E. Kaissling, J. Thorson, in Receptors for Neurotransmitters, Hormones and Pheromones in Insects, L. M. H. D. B. Sattelle, J. G. Hildebrand, Ed. (Elsevier/North-Holland, Amsterdam, 1980), pp. 261–282.

24. K. Machaca, H. C. Hartzell, Asymmetrical distribution of Ca-activated Cl channels in *Xenopus* oocytes. Biophys J 74, 1286–1295 (1998).

25. H. C. Hartzell, K. Machaca, Y. Hirayama, Effects of adenophostin-A and inositol-1,4,5-trisphosphate on Cl-currents in *Xenopus laevis* oocytes. Mol Pharmacol 51, 683–692 (1997).

26. H. C. Hartzell, Activation of different Cl currents in *Xenopus* oocytes by Ca liberated from stores and by capacitative Ca influx. J Gen Physiol 108, 157–175 (1996).

27. N. Dascal, The use of *Xenopus* oocytes for the study of ion channels. Crc Critical Reviews in Biochemistry 22, 317–387 (1987).

28. L. H. Cao et al., Odor-evoked inhibition of olfactory sensory neurons drives olfactory perception in *Drosophila*. Nat Commun 8, 1357 (2017).

29. C. Ueira-Vieira, D. A. Kimbrell, W. J. de Carvalho, W. S. Leal, Facile functional analysis of insect odorant receptors expressed in the fruit fly: validation with receptors from taxonomically distant and closely related species. Cell Mol Life Sci, (2014).

30. E. A. Hallem, M. G. Ho, J. R. Carlson, The molecular basis of odor coding in the *Drosophila* antenna. Cell 117, 965–979 (2004).

31. D. Munch, C. G. Galizia, DoOR 2.0--Comprehensive mapping of *Drosophila melanogaster* odorant responses. Sci Rep 6, 21841 (2016).

32. W. S. Leal et al., Does Zika virus infection affect mosquito response to repellents? Sci Rep 7, 42826 (2017).

## REFERENCES

1. Z. Syed, W. S. Leal, Mosquitoes smell and avoid the insect repellent DEET. Proc Natl Acad Sci U S A 105, 13598–13603 (2008).

2. W. S. Leal et al., Does Zika virus infection affect mosquito response to repellents? Sci Rep 7, 42826 (2017).

3. C. Ueira-Vieira, D. A. Kimbrell, W. J. de Carvalho, W. S. Leal, Facile functional analysis of insect odorant receptors expressed in the fruit fly: validation with receptors from taxonomically distant and closely related species. Cell Mol Life Sci 71, 4675–4680 (2014).

5. W. S. Leal, Y. M. Choo, P. Xu, C. S. da Silva, C. Ueira-Vieira, Differential expression of olfactory genes in the southern house mosquito and insights into unique odorant receptor gene isoforms. Proc Natl Acad Sci U S A 110, 18704–18709 (2013).

6. P. Xu, Y. M. Choo, A. De La Rosa, W. S. Leal, Mosquito odorant receptor for DEET and methyl jasmonate. Proc Natl Acad Sci U S A 111, 16592–16597 (2014).

7. P. Xu et al., Moth sex pheromone receptors and deceitful parapheromones. PLoS One 7, e41653 (2012).

8. P. Xu, A. M. Hooper, J. A. Pickett, W. S. Leal, Specificity determinants of the silkworm moth sex pheromone. PLoS One 7, e44190 (2012).

9. F. Zhu, P. Xu, R. M. Barbosa, Y. M. Choo, W. S. Leal, RNAi-based demonstration of direct link between specific odorant receptors and mosquito oviposition behavior. Insect Biochem Mol Biol 43, 916–923 (2013).

